# CLE4 peptide hormone regulates *de novo* shoot regeneration in potato

**DOI:** 10.1101/2024.12.06.626175

**Authors:** Maria Gancheva, Ludmila Lutova

**Affiliations:** Department of Genetics and Biotechnology, Saint Petersburg State University, Universitetskaya emb. 7/9, Saint Petersburg 199034, Russia

**Keywords:** CLE peptides, WUS, *Solanum tuberosum*, shoot regeneration, potato, CLE4

## Abstract

**Background:** *De novo* shoot regeneration involves reconstitution of stem cells under control of stem cell regulators. The WUSCHEL (WUS) transcription factor acts as a master regulator of *de novo* shoot regeneration controlling genotype-dependent regeneration efficiency. Previously we found that potato plants overexpressing the *StCLE4* gene encoding *Solanum tuberosum* CLAVATA3/EMBRYO SURROUNDING REGION-RELATED peptide hormone exhibited a *wus*-like phenotype with inhibited growth of the shoot apical meristem. We hypothesized that StCLE4 could also modulate *de novo* shoot regeneration.

**Methods and results:** In this study the role of the StCLE4 peptide in *de novo* shoot regeneration was studied by CRISPR/Cas9 mutant analysis, evaluation of overexpression effects and gene expression analysis. We found that *cle4* mutation enhanced shoot regeneration in potato.

**Conclusions:** We have found a gene in potato, the loss of which can lead to increased shoot formation. These results can be used to improve the efficiency of shoot regeneration in potato.

## 1. Introduction

*De novo* shoot regeneration refers to the process by which new shoots are formed from explants (tissues or organs) that do not contain pre-existing shoot meristems. The processes involved in shoot regeneration are complex and involve the coordination of various signaling molecules and transcription factors. Two important components that play role in shoot regeneration are the CLAVATA3/EMBRYO SURROUNDING REGION-RELATED (CLE) peptides and the WUSCHEL (WUS) transcription factor [1,2,3]. CLE peptides are small signaling peptides that have been shown to play important roles in various aspects of plant development, including shoot meristem formation and regeneration [3,4]. *Arabidopsis thaliana* CLE1–CLE7 (AtCLE1-CLE7) peptides, induced in the callus-induction medium, are actively expressed in pluripotent callus and regulate shoot regeneration [3]. Exogenous treatment with these peptides or overexpression of their precursors can inhibit shoot regeneration, whereas loss-of-function mutations in the *AtCLE1-CLE7* genes enhanced regenerative capacity [3]. The signaling mediated by the AtCLE1–CLE7 peptides requires CLAVATA1 and BARELY ANY MERISTEM receptors and leads to the repression of *WUSCHEL* (*WUS*) [3]. WUS is a homeodomain transcription factor that plays a central role in maintaining the stem cell population in the shoot meristem [5,6]. In potato (*Solanum tuberosum* L.), the WUS (StWUS) homeodomain transcription factor was recently shown to be crucial in determining regeneration capacity in different potato genotypes [7]. In genotypes with high efficiency of shoot regeneration, the expression of *StWUS* correlates with the rate of shoot regeneration, whereas *StWUS* silencing suppressed shoot regeneration [7]. At the same time, there are no data available about potato CLE (StCLE) peptides implication in plant regeneration. Previously, we found that the StCLE4 peptide is closely related to the AtCLE1-7 peptides according to phylogenetic analysis and functional studies. However, the regulation of *StCLE4* activity in potato is different from that described for the *AtCLE1-7* genes [8]. We discovered that the *StCLE4* gene is induced in potato roots under nitrogen-rich conditions and may act as a suppressor of tuberization. Plants overexpressing *StCLE4* exhibited a *wus*-like phenotype, with inhibited growth of the shoot apical meristem, and promotion of root growth and stolon conversion into branches [8]. We hypothesized that StCLE4, like AtCLE1–CLE7, could modulate *de novo* shoot regeneration.

## Materials and Methods

### Plant materials and growth conditions

The Désirée potato cultivar was obtained from the N. I. Vavilov All-Russian Institute of Plant Genetic Resources (Saint-Petersburg, Russia). This cultivar showed the high rate of shoot regeneration in the study of Park et al [7]. The potato plants were propagated on Murashige and Skoog (MS) medium with 10 g/L sucrose and 8 g/L plant agar (in a case of solid medium) in Petri dishes. Plantlets were grown under long-day conditions (16 h light/8 h dark) and a constant temperature of 24 °C.

### Constructs

For genome editing, a system of Xing et al [9] was used. To enhance the production of the Cas9 protein, the pHSE401 vector was *Xba*I-digested and the translational enhancer from japonica rice (*OsMac3*) [10,11,12,13] was introduced upstream of the *Cas9* sequence. The primer sequences used were 5′-TAA <u>tctaga</u> AAGACTAAAGAGAGCTTTTTCATAC-3′ and 5′-CCA <u>tctaga</u> ATTGCGAGACAGTGCCGTG-3′ (*Xba*I sites are underlined). The resulting construct was named pHSEe401 according to [9,14]. 19 bp *CLE4* specific sequences (5′-GACGGACCTGATCCTCGAC-3′ and 5′-AGGAAGAGGAGTACACTTA-3′) were designed to target the *StCLE4* gene using CRISPRdirect tool [15]. These sequences were cloned into the pHSEe401 vector as described previously [9]. The vector for *StCLE4* overexpression was obtained in our previous work [8]. Schematic maps of CRISPR/Cas9 and overexpression vectors were provided in Fig. S1. The resulting constructs were then used to transform *Escherichia coli* strain DH10B with calcium chloride protocol [16]. Plasmid DNA was isolated using the Plasmid Miniprep Kit (Evrogen, Moscow, Russia) and sequenced in the Research Resource Center for Molecular and Cell Technologies of Saint-Petersburg State University. For sequence and restriction sites analysis MEGA11 [17], Ugene [18], and ApE [19] were used.

Obtained vectors were introduced into *Agrobacterium tumefaciens* strain AGL1 using the freeze-thaw method [20]. *A. tumefaciens* culture was grown in YEP medium (5 g/l NaCl, 10 g/l tryptone, 10 g/l yeast extract, 15 g/l agar (in case of solid medium)) with antibiotics (40 mg/l rifampicin, 50 mg/l kanamycin). A glycerol stock of agrobacteria was stored at -80°C for future plant transformation.

### Potato plant transformation

CRISPR-induced *cle4* mutants were generated by *Agrobacterium tumefaciens*-mediated transformation mainly as described previously [21,22,23].

#### Agrobacterium tumefaciens

strain AGL1 with the required plasmid from a glycerol stock solution was cultured overnight in liquid YEP (10 ml) medium with antibiotics (40 mg/L rifampicin, 50 mg/L kanamycin) at 30ºC on a shaker (200 rpm). The next day, the culture was diluted in the antibiotic-free YEP medium (10 ml) and grown until reaching an optical density OD_600_ = 0.5-1.0. Leaf explants from plantlets were then incubated with the 150 µL agrobacterium culture in 30 ml liquid modified MS medium (mMS) (MS medium without myo-inositol, 16 g/L glucose, pH 5.8) for 15 minutes with nutation before being co-cultivated in the dark at 22°C for 2-3 days. After co-cultivation in liquid mMS medium with agrobacteria, explants were transferred onto the solid mMS medium for callus formation (CIM) (mMS with 5 mg/L IAA, 0.1 mg/L BAP and with the addition of 500 mg/L cefotaxime to prevent Agrobacterium overgrowth and 3 mg/l hygromycin for the selection of transgenic cells). The explants were cultured on this medium for 7-8 days in the 16 h light/8 h dark photoperiod and a constant temperature of 27 °C. Then, explants were transplanted onto the shoot inducing medium (SIM) – solid mMS medium with the addition of 1 mg/L BAP, 0.1 mg/L GA, 500 mg/L cefotaxime and 3 mg/L hygromycin. Every 7-14 days they were put on a fresh medium until shoots developed on the calli. The resulting 3 cm shoots were then transferred to solid MS with 10 g/l sucrose and 500 mg/L cefotaxime for rooting.

### Genotyping of edited lines

To detect editing events, total DNA of wild-type (Désirée) plants and edited lines was isolated from young leaves using the cetyltrimethylammonium bromide (CTAB) method [24]. The StCLE4 region was amplified using the ScreenMix (Evrogen, Moscow, Russia) with the following primers sequences: 5′-TGATTGTGAACCCACCTTGTC-3′ and 5′-TCAACGATCAAAAAGTGTAATGAGT-3′. The resulting fragments were isolated using a Cleanup Mini Kit (Evrogen, Moscow, Russia). PCR fragments from wild-type plants were sequenced by Sanger sequencing. PCR fragments from edited lines were cloned into the pKAN-T vector (Evrogen, Moscow, Russia) for further Sanger sequencing. Transformation of *E. coli* DH10B was performed using the standard protocol with calcium chloride [16]. Plasmid DNA was isolated using the Plasmid Miniprep Kit (Evrogen, Moscow, Russia) and sequenced in the Research Resource Center for Molecular and Cell Technologies of Saint-Petersburg State University. At least six plasmids were sequenced by Sanger sequencing for every edited line. Sanger reads were aligned to *StCLE4* wild-type sequence using the MEGA7 software [25]. Additionally, the targeted *StCLE4* region of *cle4*-cr9 and *cle4*-cr18 plants was sequenced using the Illumina MiSeq platform with MiSeq Reagent Kit v2 (2 × 250 bp). FASTQC v. 0.11.9 (http://www.bioinformatics.babraham.ac.uk/projects/fastqc/) and trimmomatic v. 0.39 programs [26] were used for the quality control and for trimming of the raw reads, respectively. Trimmed reads were mapped to a reference potato genome DM 1-3 516 R44 genome v. 6.1 using the bwa tool [27]. BAM files were sorted using samtools v. 1.13 and the visualization of data was performed using IGV [28]. Quantification and visualization of CRISPR-Cas9 results were also performed using the CRISPResso2 tool [29].

Putative off-target sites were searched in the reference potato genome SolTub_3.0 using CRISPRdirect tool [15]. These off-target regions were amplified using the Thermo Scientific Phusion high-fidelity DNA polymerase with the following primers sequences: 5′-GAGACATTTGGGTGTGTTTTGGAC-3′ and 5′-CTAGATTAAGGTGTTGGGGTTGG-3′ for first off-target region and 5′-AACCCTTCATCCACAGTTTCCT-3′, and 5′-AACAGTAACAACAAAATCTCTGGTGA-3′ for second off-target region. PCR products were sequenced in the Research Resource Center for Molecular and Cell Technologies of Saint-Petersburg State University.

### Plant regeneration assays

#### Obtaining *StCLE4*-overexpressing calli and shoots

Plant regeneration assay for evaluation of the effect of *StCLE4* gene overexpression were performed mainly as described previously [30].

Prior to transformation, the *Agrobacterium* strain AGL1 with the required plasmid was inoculated into liquid YEP medium as described above. The next day, 2 ml of the cultures were diluted in 30 ml AB-MES medium [31] with 200 µM acetosyringone, 40 mg/L rifampicin, and 50 mg/L kanamycin, and grown for several hours until reaching an optical density of 0.5-1.0 at OD_600_. The culture was then pelleted, resuspended in the infiltration medium (0.5X AB-MES components and 0.5X liquid mMS medium components with 5 mg/L IAA, 0.1 mg/L BAP and 200 µM acetosyringone), and adjusted to an optical density of 0.4-0.6 for both constructs analyzed in the experiment.

At the next stage, the leaflets were placed in falcon type tubes with an agrobacterium culture in the infiltration medium and incubated for 15 minutes with nutation. The infiltration medium was then discarded and the leaves were placed on solid mMS (with 8 g/L agar) with 5 mg/L IAA, mg/L BAP. Explants were incubated 2 days in the dark.

After cocultivation, explants were transferred onto the CIM and cultured on this medium for 8 days in the 16 h light/8 h dark photoperiod and a constant temperature of 27 °C. Then, explants were transplanted onto the SIM days at 22°C under long-day conditions. Every 7-14 days they were put on a fresh medium.

#### Obtaining *cle4* mutant calli and shoots

Hypocotyl and leaf explants of wild type and *cle4* mutants obtained due to CRISPR-Cas9 mediated gene editing were excised from plantlets and cultured on solid mMS with 5 mg/L IAA, 0.1 mg/L BAP, 500 mg/L cefotaxime for 2 or 3 days. For shoot induction, the calli were then transferred to solid mMS medium with the addition of 1 mg/L BAP, 0.1 mg/l GA, 500 mg/L cefotaxime and cultured at 22°C under long-day conditions. Every 7-14 days they were put on a fresh medium.

#### Regeneration capacity evaluation

The number of regenerated shoots from each explant was counted using a ZEISS SteREO Discovery.V12 stereomicroscope after cultivation on a SIM medium. To evaluate the statistical significance, t-test or ANOVA with further Tukey’s test were used.

#### RNA-seq Data Analysis

Raw reads were obtained from NCBI sequence read archives SRR122123 and SRR122113 (PRJNA63145 project) [32]. The quality control of the raw reads was performed using FASTQC 0.11.9 (http://www.bioinformatics.babraham.ac.uk/projects/fastqc/). The trimmomatic v. 0.39 program [26] was used to trim reads from adapters. HISAT2 v. 2.2.1 [33] was used for mapping reads to a reference potato genome DM 1-3 516 R44 genome v. 6.1. BAM files were sorted using samtools v. 1.13 [34] and the quantitation of data was performed using StringTie v. 2.2.1 [35]. Not annotated *StCLE* genes were manually included in annotation.

#### Quantitative RT–PCR Analysis

RNA isolation and reverse transcription quantitative polymerase chain reaction (RT-qPCR) were performed as previously reported [36]. For primer design, VectorNTI [37] and Ugene [18] were used. The primers were 5′-CCAGTTTTACAAGGTTGATGATTC-3′ and 5′-AGCCCACATTTACCACAATAGTG-3′ for *StUBI3* (Soltu.DM.12G001920), 5′-TGGCTAGTGCTTCTAGGTTTTTG-3′ and 5′-CGAGGATCAGGTCCGTCAGG-3′ for *StCLE4*, 5′-AAAAGAAGAGGCTCATTGCT-3′ and 5′-GATGGACACTGAACACCTGGAT-3′ for *StWUS* (Soltu.DM.02G023940). The qRT-PCR was run in three technical and three biological repeats. Student’s t-test was used to compare the difference in gene expression.

## Results

### *StCLE* genes are expressed in calli

Firstly, we analyzed expression of the *StCLE4* and *StWUS* genes in potato callus using transcriptomic data from NCBI database. Calli (10-11 week old) were generated from *S. tuberosum* group Phureja DM1-3 516 R44 leaves and stems on medium with 4.4 g/L MS salts, 30 g/L sucrose, 0.9 mg/L thiamine, 0.8 mg/L zeatin riboside, and 2 mg/L 2,4-D; 16 h light/8 h dark at room temperature [32]. Due to the fact that not all the *StCLE* genes are included in reference annotation [36], we manually included them for expression analysis. We found extremely low expression for *StCLE4* in callus and no expression for *StWUS* (Table S1). These results are consistent with previous findings that *StWUS* had low expression on CIM, and it is expressed during incubation on SIM for efficient shoot regeneration. At the same time, we found that among *StCLE* genes, *StCLE12, StCLE13*, and *StCLE17* had relatively high expression levels in calli (Table S1).

### *StCLE4* downregulation improves shoot regeneration

To verify whether StCLE4 is involved in shoot regeneration in potato, we generated plants with *StCLE4* precursor overexpression and CRISPR/Cas9-induced *cle4* mutants. For *StCLE4* overexpression, we used a vector construction obtained previously [8]. We transformed leaf explants with constructions for overexpression of *StCLE4* or with control construction for *GFP* overexpression (kindly provided by Dr Tvorogova from Saint Petersburg State University). After 75 days of cultivation, shoots were regenerated from transgenic calli, and we estimated the number of shoots per callus. We found no significant differences between the calli transformed with the construction for *StCLE4* overexpression and the control calli (Fig. 1a).

**Fig. 1.**
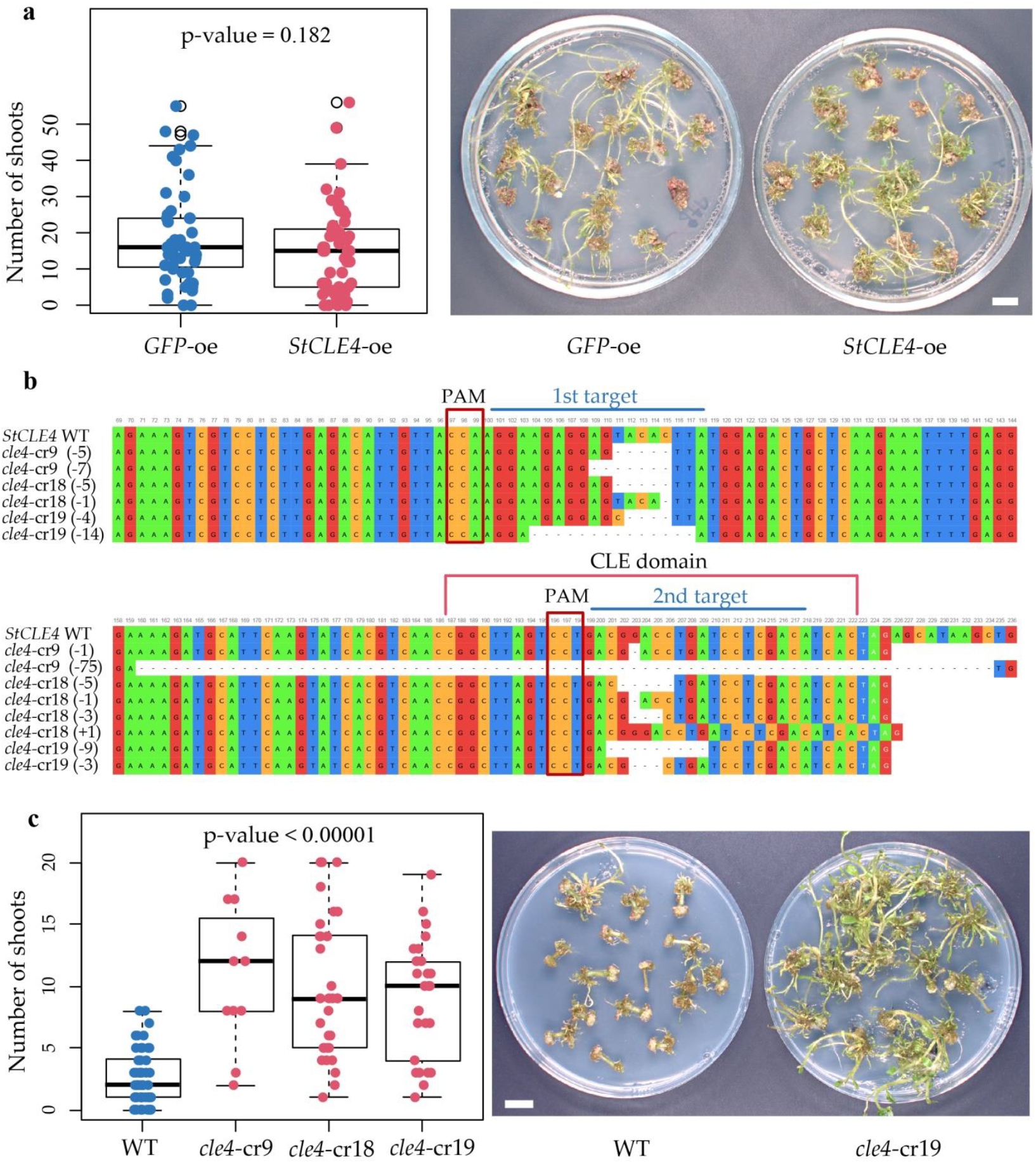
Effects of the *StCLE4* gene overexpression and loss-of-function on the *de novo* shoot regeneration. **a** Shoot regeneration of explants transformed with the construction for *StCLE4* (*StCLE4*-oe) and *GFP* (*GFP*-oe) overexpression. The Student’s t-test is used to compare the means between two groups. **b** DNA sequences of edited (*cle4-*cr9, *cle4-*cr18, *cle4-*cr19) and wildtype (WT) alleles of the *StCLE4* gene. Only regions with targets are displayed. The lengths of deletions (-) or insertions (+) are given in brackets. Stop codon is marked in white. The default coloring in the Ugene [18] is used. **c** Shoot regeneration of wild type Désirée plants (WT) and CRISPR/Cas9-induced *cle4* mutants. p-value indicates statistically significant differences compared with WT as determined by ANOVA with further Tukey’s test. Boxplot were generated using boxplot package in R. Scale bar = 1 cm.

We next characterized the function of the CLE4 peptide in *de novo* shoot regeneration by analyzing loss-of-function mutants that were generated using the CRISPR/Cas9 genome engineering technology. We developed construction with the translational enhancer *OsMac3* upstream of the *Cas9* gene in the pHSEe401 vector and targeted two different sites in the *StCLE4* gene that facilitate genome-editing events in polyploid potato [12] (Fig S1). Twenty regenerants (named *cle4*-cr1-20) were obtained. Sequencing has shown that *cle4*-cr9, *cle4*-cr18, and *cle4*-cr19 mutants contained deletions and insertions of different lengths (Fig. 1b, Fig. S2). Wild-type alleles were detected only in 4.9% of reads in the second target in the *cle4-cr*18 plant genome (Fig. S2). We also analyzed possible off-targets predicted by the CRISPRdirect program [15]. Off-targets are located in noncoding regions of the *ULTRAPETALA2* (PGSC0003DMG400043421) and *SWI/SNF-RELATED MATRIX-ASSOCIATED ACTIN-DEPENDENT REGULATOR OF CHROMATIN SUBFAMILY-*RELATED (PGSC0003DMG400040275) genes. We did not find any changes in these sequences, except that the Désirée has an insertion of three nucleotides in the off-target region (Fig. S3).

We analyzed regeneration capacity of both hypocotyl and leaf explants after 38 days of incubation explants on a SIM and found that *cle4* mutant calli formed more shoots than control calli (Fig. 1c, Fig. S4). We supposed that StCLE4 could limit *de novo* shoot regeneration capacity of potato explants. Since WUS is a critical transcription factor for shoot regeneration in *Arabidopsis* [38] and potato [7], we evaluated its expression in hypocotyl explants after 12 days incubation on a SIM. We found no activation of *StWUS* in *cle4* mutant calli (Fig. S5).

## Discussion

Shoot apical meristems develop from explant cells by *de novo* shoot regeneration mechanism. The first stage involves reprogramming of differentiated cells and callus formation. In this study we analyzed transcriptomic data of 10 and 11 week old potato calli that were incubated in callus induction media and found that *StCLE12, StCLE13*, and *StCLE17* had relatively high expression levels (Table S1), but there are no studies on the role of these genes in calli development. StCLE12 participates in the regulation of vascular cell proliferation, leaf senescence and tuber development in potato [39]. StCLE13 is grouped with AtCLE46 that homologous to AtCLE41 and AtCLE44, playing crucial roles in vascular meristem maintenance [40,41]. StCLE17 is close to AtCLE13, which promoter is active in root cap and root apical meristem [42]. At the same time, previous studies of the shoot regeneration of Arabidopsis have shown that calli induced by incubating explants on callus induction media had many of the characteristics of lateral root primordia and root apical meristem [43,44]. It is also known, that the AtCLE41/44 peptides regulate lateral root formation by modulating the auxin transport [45]. We speculated that together with root-expressed AtCLE13 they may affect callus formation.

At the next stage of shoot regeneration, *de novo* shoot apical meristems should be developed in pluripotent callus. The AtCLE1–7 peptides are involved in this process [3]. These peptides modulate shoot regeneration capacity by repression of *AtWUS* expression, which is essential for stem cell induction and shoot apical meristem initiation during shoot regeneration. In potato, high *StWUS* expression level is also essential for efficient shoot regeneration [7]. Previously we found that the *StCLE4* gene belongs to the same clade of *CLE*s as *AtCLE1-7* [36], and plants overexpressing *StCLE4* demonstrated a *wus*-like phenotype, with the arrested growth of the shoot apical meristem [8]. However, opposite to *A. thaliana CLE1, -3, -4*, and *-7* genes, the *StCLE4* gene expression was significantly induced by nitrogen [36]. Previously, it was shown that nitrogen nutrition is necessary for shoot organogenesis [46,47,48], and we hypothesized that the nitrogen-activated *StCLE4* gene may be involved in shoot regeneration as positive regulator. Unexpectedly, we found that shoot regeneration in explants transformed with the construction for *StCLE4* overexpression was not affected, whereas loss of its function enhanced shoot regeneration. We also found that *cle4* mutation did not significantly alter *StWUS* expression in stem explants after 12 days of incubation on a SIM. It is possible that changes in *StWUS* expression may occur at other time points. In addition, it can also be speculated that StCLE4-mediated signaling pathway may suppress shoot regeneration by regulating other genes involved in this process. Further studies on the mechanisms of shoot regeneration regulation by CLE peptides and the role of external factors, such as nitrogen, in this process are needed.

## Conclusion

Thus, we have discovered a gene in potato, the loss of which can lead to an increased shoot formation. These results can be used to enhance shoot regeneration capacity of low-efficiency potato genotypes.

## Supplementary Information

Fig. S1. Schematic maps of CRISPR/Cas9 and overexpression vectors.

Fig. S2. Alelles around cut sites in *cle4*-cr9 and *cle4*-cr18 mutants quantified and visualized by CRISPResso2.

Fig. S3. PCR genotyping of off-target Cas9 activity.

Fig. S4. Shoot regeneration of wild type Désirée plants (WT) and CRISPR/Cas9-induced *cle4* mutants (*cle4*-cr) after 32 days of incubation of leaf explants on a SIM.

Fig. S5. Expression of *StWUS* and *StCLE4* in hypocotyl explants of wild type Désirée plants (WT) and CRISPR/Cas9-induced *cle4* mutants (*cle4*-cr) after 12 days incubation on a SIM.

Table S1. Expression profiles of the *StCLE* and *StWOX* genes in calli.

## Statements & Declarations

## Author Contributions

Maria Gancheva designed and performed the experiments, collected data, prepared the figures, and wrote the article with revisions from Ludmila Lutova. All authors have read and agreed to the published version of the manuscript.

## Data Availability

Raw reads of the *StCLE4* region of *cle4*-cr9 and *cle4*-cr18 plants have been deposited at the National Center for Biotechnology Information database under SRA numbers <u>SRR30481987</u> and <u>SRR30481988</u> (BioProject number <u>PRJNA1154414)</u>.

## Acknowledgments

We are grateful to Varvara Tvorogova and Maria Lebedeva (Department of Genetics and Biotechnology, Saint Petersburg State University) for fruitful discussions and the critical reading of the manuscript; and the Research Resource Center for Molecular and Cell Technologies of Saint-Petersburg State University for DNA sequencing.

