## Supplemental Figures and Table for "CLE4 peptide hormone regulates *de novo* shoot regeneration in potato"

Fig. S1. Schematic maps of CRISPR/Cas9 and overexpression vectors. **a** The StCLE4 region sequence with target DNA regions and primers used for genotyping (blue). The coding region of the *StCLE4* gene is highlighted in green. **b** pHSEe401 vector for CRISPR/Cas9 editing and pMDC32 vector for overexpression. RB—right T-DNA border; gRNA—guide RNA; NLS—nuclear localization signal; Cas9—CRISPR associated nuclease 9; LB—left T-DNA border.

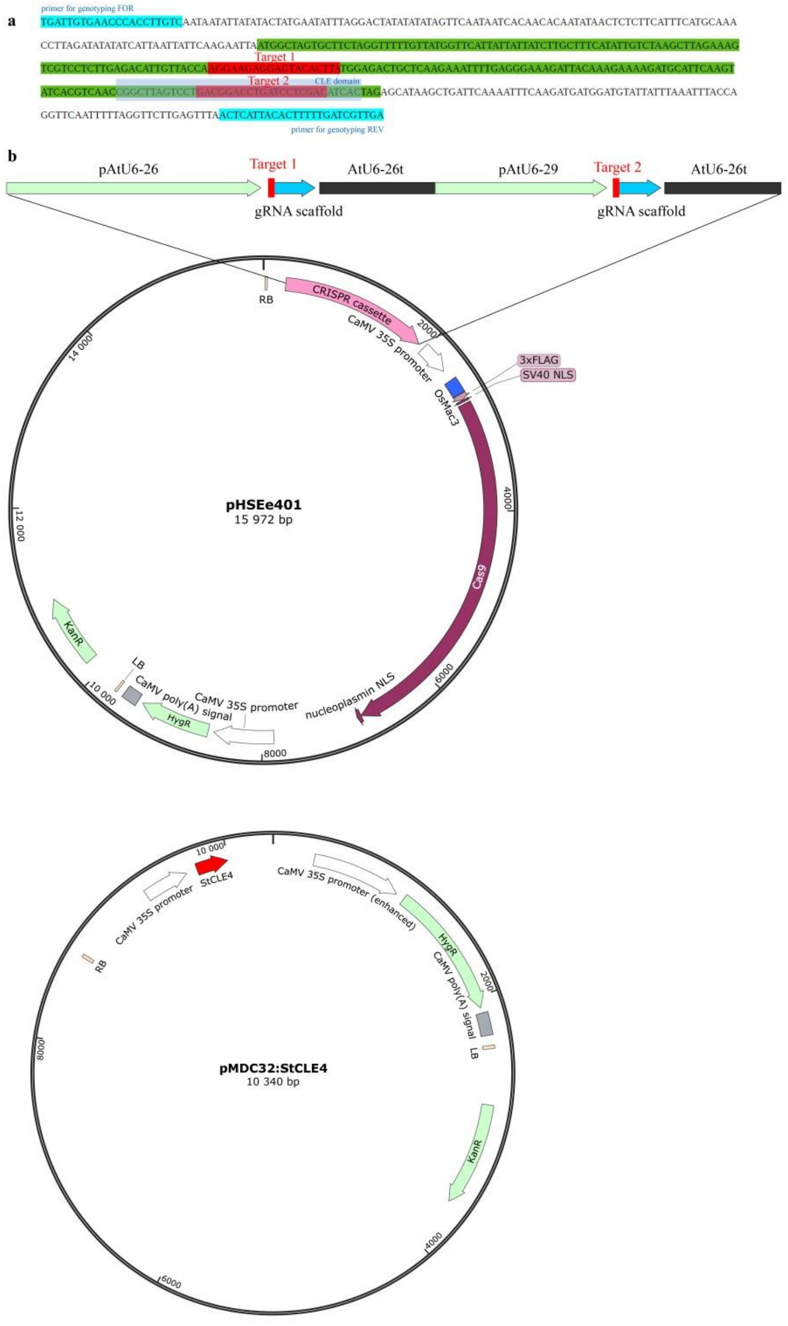

Fig. S2. Alelles around cut sites in *cle4*-cr9 and *cle4*-cr18 mutants quantified and visualized by CRISPResso2. Alleles occurring in less than 3% of reads were removed.

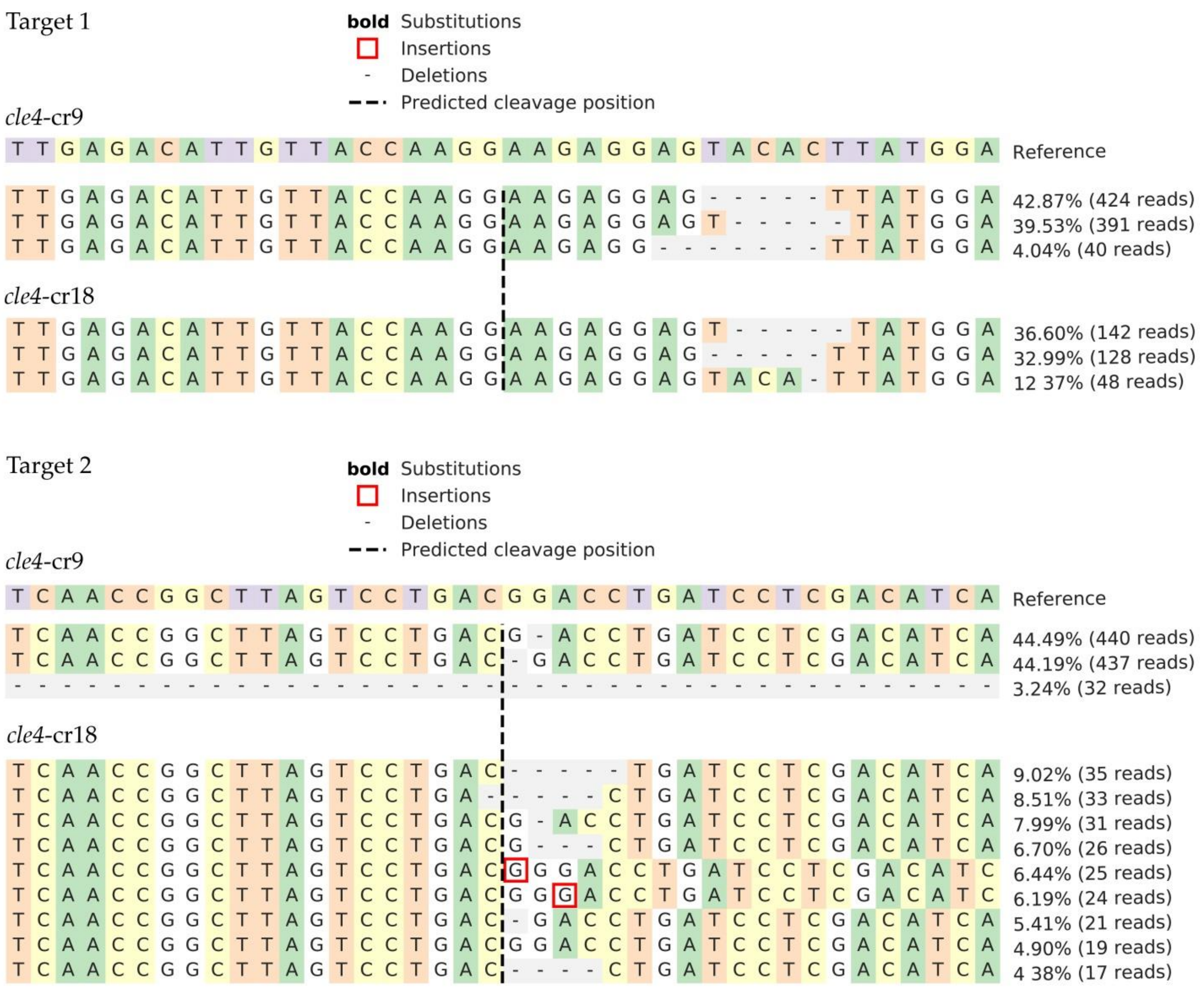


Fig. S3. PCR genotyping of off-target Cas9 activity.

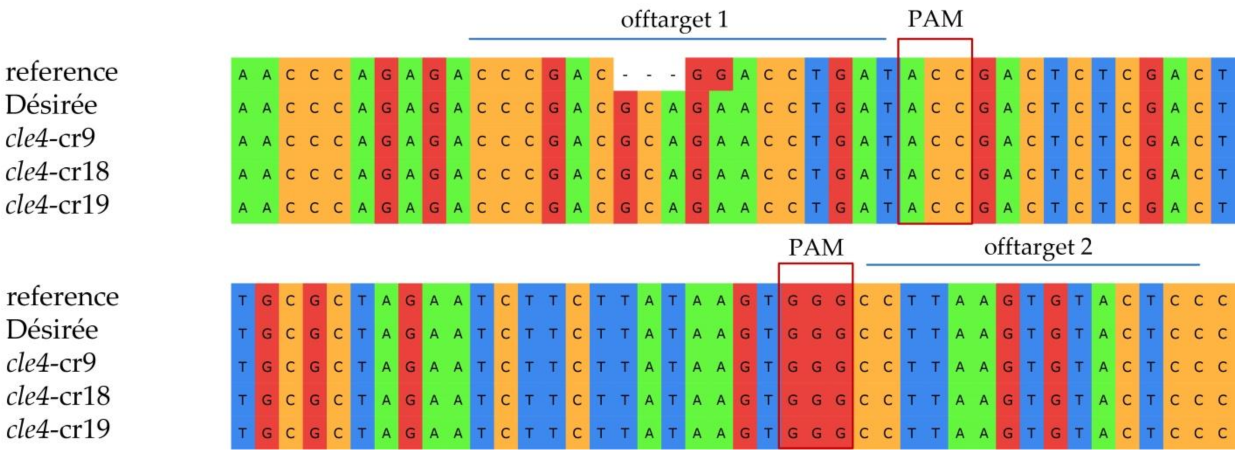

Fig. S4. Shoot regeneration of wild type Désirée plants (WT) and CRISPR/Cas9-induced *cle4* mutants (*cle4*-cr) after 32 days of incubation of leaf explants on a SIM. Boxplots were generated using boxplot package in R. Scale bar = 2 mm.

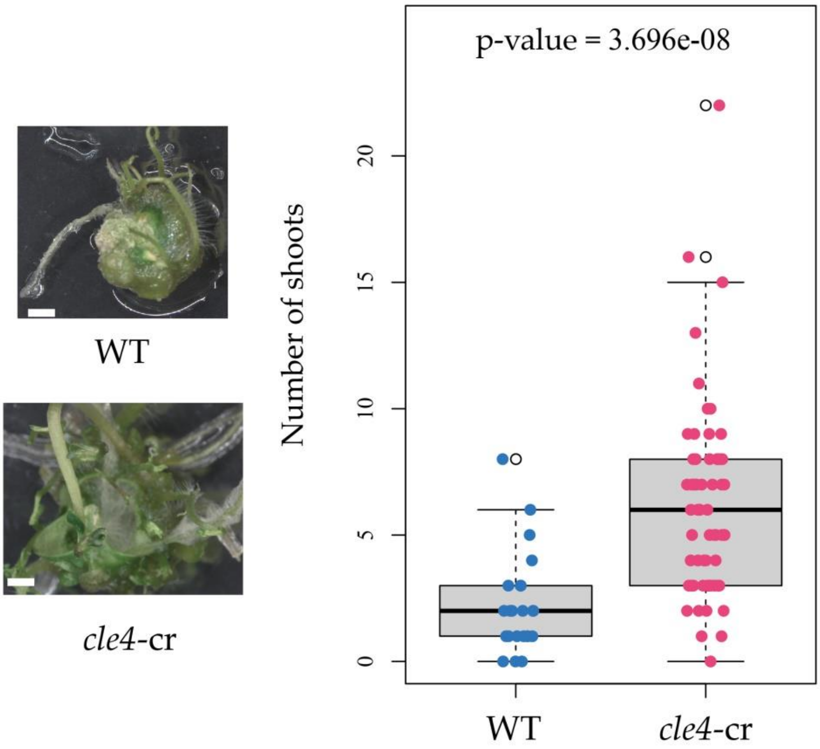


Fig. S5. Expression of *StWUS* and *StCLE4* in hypocotyl explants of wild type Désirée plants and CRISPR/Cas9-induced *cle4* mutants (*cle4*-cr) after 12 days incubation on a SIM. The gene expression levels were normalized to 1 against the expression found in the *cle4*-cr19 plants. Boxplots were generated using boxplot package in R. Scale bar = 2 mm.

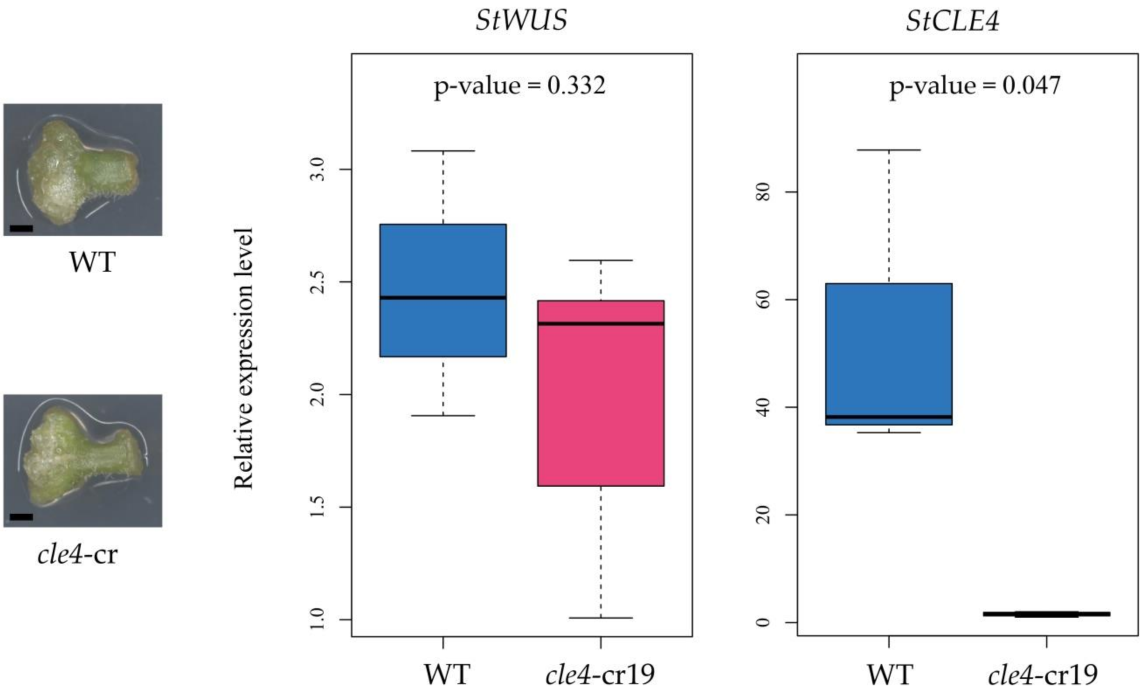

Supplementary Table 1. Expression profiles of *StCLE* and *StWOX* genes in calli.

| | SRR122113_gene_abundance | | | SRR122123_gene_abundance | | | |
|---|---|---|---|---|---|---|---|
| Gene Name | Coverage | FPKM | TPM | Coverage | FPKM | TPM | Gene ID |
| *StCLE1* | 8 | 5 | 9 | 2 | 5 | 7 | |
| *StCLE10* | 0 | 0 | 0 | 0 | 0 | 0 | |
| *StCLE11* | 0 | 0 | 0 | 2 | 6 | 9 | |
| *StCLE12* | 38 | 23 | 44 | 15 | 39 | 56 | |
| *StCLE13* | 81 | 50 | 93 | 38 | 97 | 139 | |
| *StCLE14* | 0 | 0 | 0 | 0 | 0 | 0 | |
| *StCLE15* | 0 | 0 | 0 | 0 | 0 | 0 | |
| *StCLE17* | 20 | 12 | 23 | 11 | 28 | 40 | |
| *StCLE18* | 0 | 0 | 0 | 0 | 0 | 0 | |
| *StCLE2* | 0 | 0 | 0 | 0 | 0 | 0 | |
| *StCLE20* | 0 | 0 | 1 | 0 | 0 | 0 | |
| *StCLE22* | 3 | 2 | 3 | 1 | 4 | 5 | |
| *StCLE23* | 0 | 0 | 0 | 0 | 0 | 0 | |
| *StCLE24* | 0 | 0 | 0 | 0 | 0 | 0 | |
| *StCLE25* | 0 | 0 | 0 | 0 | 0 | 0 | |
| *StCLE26.2* | 0 | 0 | 0 | 0 | 0 | 0 | |
| *StCLE27* | 0 | 0 | 0 | 0 | 0 | 0 | |
| *StCLE28* | 0 | 0 | 0 | 1 | 1 | 2 | |
| *StCLE29* | 0 | 0 | 0 | 0 | 0 | 0 | |
| *StCLE30* | 0 | 0 | 0 | 0 | 0 | 0 | |
| *StCLE31* | 0 | 0 | 0 | 0 | 0 | 0 | |
| *StCLE32* | 1 | 1 | 1 | 1 | 1 | 2 | |
| *StCLE33* | 0 | 0 | 0 | 0 | 0 | 0 | |
| *StCLE34* | 1 | 0 | 1 | 0 | 0 | 0 | |
| *StCLE35* | 0 | 0 | 0 | 0 | 0 | 0 | |
| *StCLE36* | 0 | 0 | 0 | 0 | 0 | 0 | |
| *StCLE38* | 0 | 0 | 1 | 0 | 0 | 0 | |
| *StCLE39* | 0 | 0 | 0 | 0 | 1 | 2 | |
| *StCLE4* | 1 | 0 | 1 | 1 | 2 | 3 | |
| *StCLE40.1* | 0 | 0 | 0 | 0 | 0 | 0 | |
| *StCLE41* | 0 | 0 | 0 | 0 | 0 | 0 | |
| *StCLE42* | 0 | 0 | 0 | 0 | 0 | 0 | |
| *StCLE5* | 0 | 0 | 0 | 0 | 0 | 0 | |
| *StCLE7* | 2 | 1 | 2 | 1 | 2 | 2 | |
| *StCLE9* | 2 | 1 | 2 | 0 | 1 | 1 | |
| *StWOX1* | 0 | 0 | 0 | 0 | 0 | 0 | Soltu.DM.03G033140 |
| *StWOX11* | 3 | 2 | 4 | 1 | 2 | 3 | Soltu.DM.06G028250 |
| *StWOX13* | 107 | 66 | 124 | 34 | 86 | 124 | Soltu.DM.02G022590 |
| *StWOX14* | 0 | 0 | 0 | 0 | 0 | 0 | Soltu.DM.11G026210 |
| *StWOX2* | 0 | 0 | 0 | 0 | 0 | 0 | Soltu.DM.06G031390 |
| *StWOX3a* | 0 | 0 | 0 | 0 | 0 | 0 | Soltu.DM.11G026200 |
| *StWOX3b* | 0 | 0 | 0 | 0 | 0 | 0 | Soltu.DM.06G031020 |
| *StWOX4* | 29 | 18 | 34 | 8 | 21 | 31 | Soltu.DM.04G033400 |
| *StWOX5* | 18 | 11 | 21 | 5 | 12 | 18 | Soltu.DM.03G012800 |
| *StWOX9* | 0 | 0 | 0 | 0 | 0 | 0 | Soltu.DM.02G016680 |
| *StWUS* | 0 | 0 | 0 | 0 | 0 | 0 | Soltu.DM.02G023940 |
